# Uniaxial loading induces a scalable switch in cortical actomyosin flow polarization and reveals mechanosensitive regulation of cytokinesis

**DOI:** 10.1101/560433

**Authors:** Deepika Singh, Devang Odedra, Christian Pohl

## Abstract

During animal development, it is crucial that cells can sense and adapt to mechanical forces from their environment. Ultimately, these forces are transduced through the actomyosin cortex. How the cortex can simultaneously respond to and create forces during cytokinesis is not well understood. Here we show that under mechanical stress, cortical actomyosin flow switches its polarization during cytokinesis in the *C. elegans* embryo. In unstressed embryos, longitudinal cortical flows contribute to contractile ring formation, while rotational cortical flow is additionally induced in uniaxially loaded embryos. Rotational cortical flow is required for the redistribution of the actomyosin cortex in loaded embryos. Rupture of longitudinally aligned cortical fibers during cortex rotation releases tension, initiates orthogonal longitudinal flow and thereby contributes to furrowing in loaded embryos. A targeted screen for factors required for rotational flow revealed that actomyosin regulators involved in RhoA regulation, cortical polarity and chirality are all required for rotational flow and become essential for cytokinesis under mechanical stress. In sum, our findings extend the current framework of mechanical stress response during cell division and show scaling of orthogonal cortical flows to the amount of mechanical stress.

## 1. Introduction

While cells remodel their actomyosin cortex during cell division, they have to simultaneously integrate chemical and mechanical stimuli from the local environment to ensure successful cytokinesis. In order for cytokinesis to be robust yet responsive to extrinsic stimuli, three fundamental control principles have evolved, (a) redundancy [1], (b) mechanosensitivity [2], and (c) positive/negative feedback [3]. Examples for these control principles are (a) partially redundant molecular motors, actin cross-linkers, and membrane trafficking pathways, (b) molecular mechanosensitivity of integral cytokinesis proteins such as non-muscle myosin II, α-actinin, and filamin [4, 5], and (c) RhoA-dependent self-enhancing local assembly and contraction of actomyosin as well as astral microtubule-based suppression of actomyosin contractility [6], which both are required to generate cortical contractile actomyosin flow during cell division.

Work in the last decade has led to the identification of the main mechanosensory system that operates during cell division. The core of this system is non-muscle myosin II, which can amplify sensed forces through its lever arm [4], and which shows mechanosensitive accumulation through cooperative binding to F-actin [7]. This results in a positive feedback on the assembly of non-muscle myosin II bipolar thick filaments [2]. In addition, for other mechanosensitive proteins, two conserved and distinct modes of force-dependent accumulation have been recently demonstrated, a rapid, diffusion-based mode due to tensile forces increasing the lifetime of the F-actin bound state (catch bonding), and a slower mode due to non-muscle myosin-II-dependent cortical flow [5]. The latter serves as an additional biomechanical positive feedback, strongly suggesting that cytokinesis control principles operate interdependently.

Among the control principles mentioned above, feedback during cytokinesis crucially depends on spindle microtubules since they constitute key modulators of cortical contractility [3]. Wolpert’s and Rappaport’s classical experiments have led to the astral relaxation model in which astral microtubules soften the polar cortex (by suppressing actomyosin contractility) while the equatorial cortex stiffens during division. Very recently, it was shown that polar clearing of contractile ring components requires TPXL-1-dependent cortical activation of Aurora A [6].

Moreover, the ability of the actomyosin cortex to contract and to generate long-range flow not only depends on the non-muscle myosin II motor protein but also on the spatial organization of actin filaments (polarization, branching, bundling) and their connectivity (degree and density of crosslinks) [8]. Recently, it has been shown that non-muscle myosin II-powered cortical actomoysin flow leads to contractile ring formation by alignment of actin filaments in the *C. elegans* one cell embryo due to compression of the gel-like cortex in the equatorial region [9]. This suggests that non-muscle myosin II-dependent flow indeed re-organizes the cortical actin network during cytokinesis as has been proposed previously [10]. It also suggests that self-enhancing feedback mechanisms are generally involved in self-organization of the cytokinetic cortex and involved particularly in forming the contractile ring. However, furrow formation due to coupling of cortical flow and actin alignment apparently only enhances but is not required for cytokinetic ring formation [9]. Moreover, actomyosin dynamics and architecture as well as cortical contractile actomyosin flows seem to variably contribute to cytokinesis progression when comparing different systems [3, 9, 11–17].

Furthermore, cortical contractile actomyosin flows in the *C. elegans* embryo are strictly dependent on RhoA activation and do not only cause translation of the cortex (like during anteroposterior polarization) [18] but also its rotation immediately before division of the two-cell embryo [19, 20] and during chiral symmetry breaking [21, 22]. Importantly, whole cell cortex rotation occurs during cell division when cytokinetic actomyosin nodes have formed. This mesoscopic rotational flow is most likely due to generation of torque at the molecular level: showed using in vitro assays that during myosin-driven sliding of actin filaments, a torque component can be observed [23] that induces a right-handed rotation of an actin filament around its long axis with one revolution per sliding distance of approximately 1 μm [24]. Similar rotation or twirling of actin filaments have been confirmed in more recent reports [25, 26]. Although the molecular origin of torque in actomyosin dynamics is well understood, how torque leads to coordinated cortical rotational dynamics remains unexplored.

The division of the one-cell *C. elegans* embryo represents a highly suitable model to quantitatively dissect spatiotemporal dynamics of the cytokinetic actomyosin cortex and to uncover underlying regulatory principles [9, 18, 27–33]. Previously, it has been shown through highly informative ablation experiments of the contractile ring that it is able to repair requiring an increased tension in the ring and reduced cortical tension in the vicinity [34]. This strongly suggests that global cortical dynamics respond to mechanical stress during cytokinesis that might require differential regulation of cytokinetic cortical flow. Here, we quantitatively describe the biomechanical responses to a different type of stress, loading. For this, progressive uniaxial compression is used in the form of the classical parallel plate assay [35–37]. With this mechanical manipulation, it is possible to demonstrate that a recently uncovered type of polarizing cortical flow, rotational flow [19–21], is mechanoresponsive, scales to the amount of load and contributes to successful division when cells experience mechanical stress. Anisotropic mechanosensitive accumulation of non-muscle myosin II suggests that cortical stress is similarly anisotropic in uniaxially loaded embryos as has been recently shown for uniaxially loaded mammalian cells [38]. Importantly, rotational flow leads to a re-arrangement of the anisotropically distributed of actomyosin in loaded embryos. Cortical rotation requires a broad set of actomyosin regulators of which several only become essential for cytokinesis under mechanical stress. Hence, our data suggests that the main biological role of cortical flow repolarization during cytokinesis lies in balancing spatial and tension anisotropies in the cortex and that converging longitudinal flow is required for successful furrowing in mechanically stressed embryos.

## 2. Materials and Methods

### 2.1 Worm Strains, maintenance and RNA interference

Integrated *C. elegans* strains expressing lifeact-fusion proteins expressed from pie-1 promoters have been described elsewhere [39, 40]. Strains JJ1473 (zuIs45), LP162 (*nmy-2(cp13*), and RW10223 (itIs37; stIs10226) were provided by the Caenorhabditis Genetics Center (CGC), which is funded by NIH Office of Research Infrastructure Programs (P40 OD010440). Strains were maintained under standard conditions [41]. RNAi was performed by feeding using clones from commercially available libraries [42, 43].

### 2.2 Microscopy and laser ablation

Embryo preparation and mounting has been described elsewhere [39, 44]. Mounting was modified by using differently sized polystyrene (15μm, 20μm, 25μm; Polysciences, Hirschberg, Germany) and polymethylmethacrylate spheres (12μm and 13.5μm, PolyAn, Berlin, Germany). Microscopy was performed with a VisiScope spinning disk confocal microscope system (Visitron Systems, Puchheim, Germany) based on a Leica DMI6000B inverted microscope, a Yokogawa CSU X1 scan head, and a Hamamatsu ImagEM EM-CCD. All acquisitions were performed at 21°C–23°C using a Leica HC PL APO 63×/1.4-0.6 oil objective. Cell cortex ablations were performed using a pulsed 355 nm UV laser mounted on the same microscope. One ablation cycle was performed per acquisition with a residence time per pixel of 3.5 ms. Acquisitions pre-and post-ablation were performed with 200 ms intervals.

### 2.3 Particle image velocimetry (PIV)

PIV analysis was performed on maximum intensity projected images using a custom version of PIVlab developed for MATLAB [44, 45]. This customized software is available from the authors upon request. Specifically, two pass interrogation windows of 64×64 pixels and 32×32 pixels with 50% overlap were used to map consecutive frames acquired at 2 s intervals. To align the biological time of flow across embryos, we choose foci formation as starting point. To calculate vector maps, correlation between subsequent windows was computed using fast Fourier transformation (FFT).

### 2.4 Quantification and kymograph representation of flow profiles

The flow profile for each time point was projected on the long axis of the embryo by dividing the whole vector profile of the embryo into 13 bins and taking a mean along the short axis. A time course profile or kymograph was obtained by averaging bin velocities for 5 embryos in each condition. For visualization, heat maps were generated after applying cubic interpolation using a custom MATLAB script. Variability between embryos for each condition was estimated by calculating standard error of mean.

### 2.5 Measurements

NMY-2 and TBB-2 signal intensities, NMY-2 node number and size, NMY-2 filament contraction rate of linearly organized NMY-2, cortical residence times of NMY-2 and lifeact, NMY-2 outward flow velocities, spindle microtubule angles, as well as furrow asymmetry and anterior-posterior domain sizes were manually measured in ImageJ using the built-in toolset. Cortical residence times were measured from traces in kymographs or by tracking cortical structures in sequential frames of high-resolution time lapse series. Longitudinal flow range was measured in the anterior domain by extracting continuous tracks from PIV data that show velocities higher than 0.5 μm/min and normalizing them to embryo length. Cleavage success was manually quantified by inspecting time-lapse microscopy data.

For shape parameter quantification of embryos in utero (Figure 3A), the embryo perimeter was segmented using a custom MATLAB script by applying a median filter and thresholding. Circularity was defined as 4π(area/permieter^2^).

Calculation of curvature to quantify blebbing (Figure 3G) was performed by segmentation of cell boundaries using a custom MATLAB script. For each time point, the boundary at the anterior end of the embryo was divided into 400 equidistant points. A circle was fit for each boundary point using this point and two boundary points that were four points away. The local curvature was defined as reciprocal of the radius of this fitted circle.

To establish that the *C. elegans* embryo follows Laplace’s law (Figure 2A), sideview projections of embryos were obtained by using a custom MATLAB script. Projected images were denoised (Wiener filter) and the embryo’s boundary was segmented by adaptive thresholding. For each point on the boundary, a circle was fitted on three points with a spacing of 30 points. Curvature was defined as the inverse of the radius of the fitted circle. Contact angles were measured based on segmented boundaries.

## 3. Results

### 3.1 Convergent longitudinal flow polarizes cortical NMY-2

In order to establish an unbiased readout for cortical dynamics during cytokinesis, we performed time-lapse microscopy with high spatiotemporal resolution of the first division in wild type (wt) *C. elegans* embryos expressing NMY-2::GFP (a CRISPR/Cas9 edited GFP-fusion of an essential non-muscle myosin II gene) [46]. This data (Figure 1A) was then subjected to quantitative analysis by particle image velocimetry (PIV). PIV tracking revealed longitudinal cortical NMY-2 flows with opposite direction, from anterior (6±0.05 μm/min, n = 5) and posterior poles (6.5±0.09 μm/min, n = 5) towards the cell equator (Figure 1B, top panel; Video 1). Convergence of these flows at the equator leads to the transformation of cortical NMY-2 nodes (1.8±0.1 μm in diameter, n = 25) into parallel, linearly organized NMY-2 (0.25-0.5 μm in width and 3.5±0.6 μm in length, n = 20; Figure 1A,C), which first form a narrow stripe (6.8±0.07 μm; n = 5) that subsequently becomes part of the incipient contractile ring by alignment and bundling (Figure 1D; Video 1).

**Figure 1.**
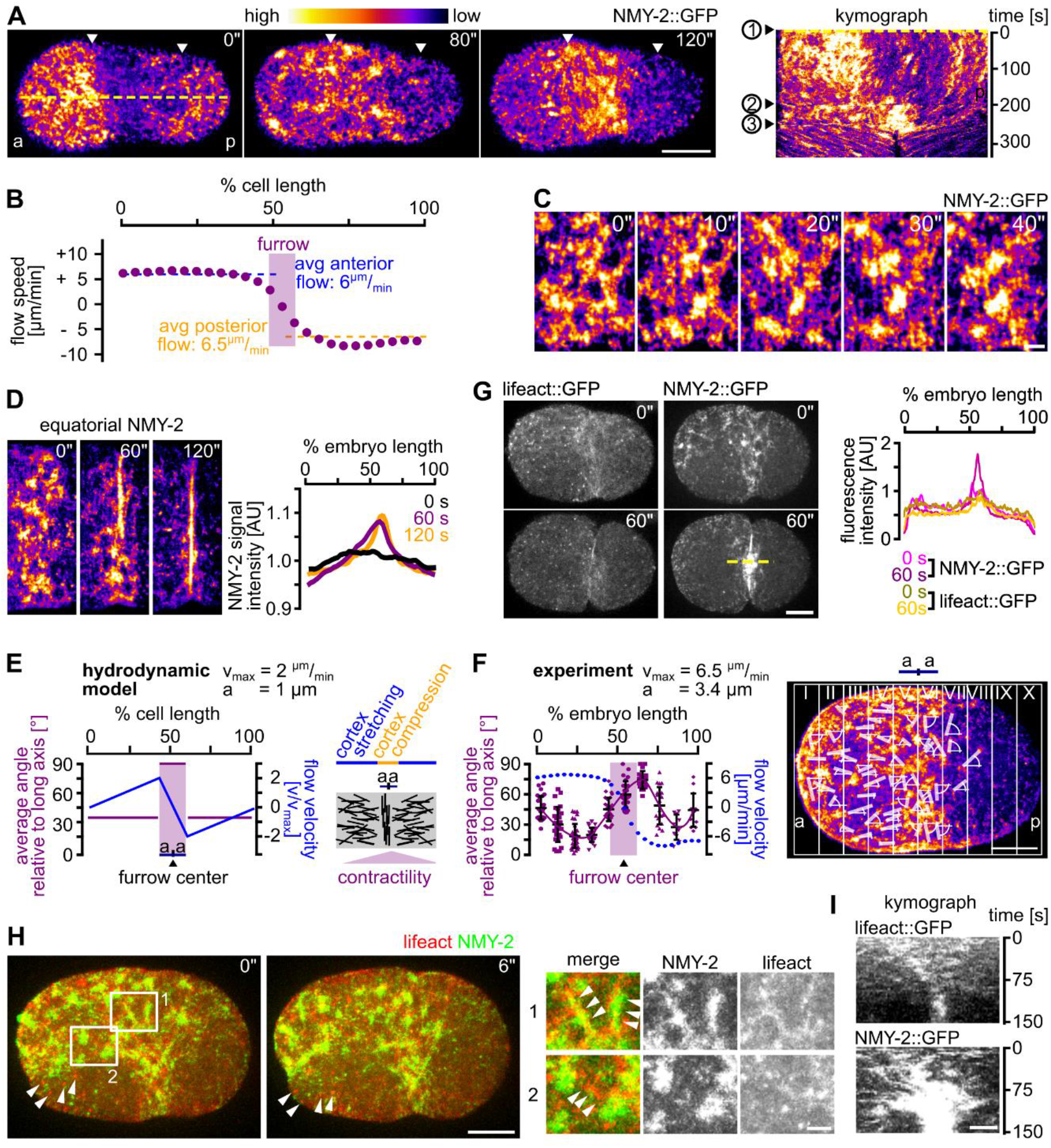
Longitudinal flow is organizes cortical NMY-2 during contractile ring formation. **(A)** Left: Maximum projected stills from time lapse microscopy of embryos expressing NMY-2::GFP. White arrow heads mark the boundaries of the anterior and posterior NMY-2 caps upon polarization. Right: Kymograph along the yellow dashed line in the leftmost panel. Numbers on the left refer to onset of cap formation (1), onset of NMY-2 cytokinetic foci formation (2), and start of furrow invagination (3). Scale bar = 10 μm. See also Video 1. **(B)** Average cortical NMY-2 flow velocity profile along the a-p axis generated from PIV data of 5 embryos over the time window of longitudinal flow (60 s). **(C)** Maximum projected stills from time lapse microscopy of the furrow region; scale bar = 2.5 μm. **(D)** Left: Stills from maximum projected embryos showing NMY-2 dynamics at the equatorial ring. Scale bar = 2.5 μm. Right: Normalized NMY-2::GFP signal intensities along the a-p axis in one-cell embryos. Intensity profiles at 0 s, 60 s and 120 s are represented by black, purple and yellow traces, respectively (n = 5 each). **(E)** Quantification of NMY-2 linear orientation. Left: Distribution of order parameter and flow velocity for a cylindrical system undergoing cytokinesis (see cartoon according to [47]). **(F)** Left: Measured angle and flow velocities along the a-p axis (n = 5). Right: Representative embryo with angles of linearly organized NMY-2 relative to the a-p-axis. **(G)** Left: Maximum projected stills from time lapse microscopy of representative wt embryos expressing either lifeact::mCherry or NMY-2::GFP. Scale bar = 10 μm. Right: Quantification of signal intensities from the embryos depicted in the middle panel. **(H)** Left: Organization of NMY-2 and F-actin during onset of cytokinesis. Maximum projected stills from time lapse microscopy of embryos expressing lifeact::mCherry and NMY-2::GFP. White arrow heads mark persistent actin foci. Scale bar = 10 μm. Right: Enlarged cortical areas from left panels showing localization of NMY-2 on actin filaments (1) and NMY-2 foci connected by actin filaments (2). Scale bar = 2.5 μm. **(I)** Kymographs for lifeact (top) and NMY-2 (bottom) at the midbody region (generated along the dashed yellow line in panel G). Scale bar = 2.5 μm.

Previously, a physical model based on hydrodynamic active gel theory has explained formation of the F-actin component of the contractile ring by cortical flow [9, 47]. In this model, opposing flows that emerge at the poles and converge at the equator promote ordering of cortical actin filaments into parallel bundles (Figure 1E). In agreement with the model, our analyses revealed similar flow velocity profiles for NMY-2 and similar ordering during cortical flow (Figure 1F). Therefore, cortical flow not only polarizes F-actin but also NMY-2, thereby promoting contractile ring formation form linearly organized NMY-2 that undergoes bundling in the equatorial region (Figure 1E) [9]. Hence, hydrodynamic active gel theory combined with PIV-based NMY-2 cortical flow analysis seem well suited to investigate cytokinesis mechanics.

While analysis of NMY-2 foci dynamics during longitudinal flow revealed an average lifetime of 29±2 s (n = 25), analysis of F-actin (using lifeact::mCherry; [40]) shows that it does not concentrate in cortical NMY-2 foci and forms much smaller, uniformly sized (0.4±0.1 μm; n = 20) and long-lived (124±48 s; n = 20) foci that do not undergo changes during cytokinesis (Figure 1G). Nevertheless, NMY-2 decorates actin filaments shortly after onset of cytokinesis; while actin filaments disassemble subsequently after around 15 s, linearly organized NMY-2 and NMY-2 foci have substantially longer half-lives (Figure 1H). Consistent with F-actin showing faster cortical turnover, we also find that F-actin shows slightly weaker longitudinal flow with a shorter range (0.3±0.2 embryo lengths) when compared to NMY-2 (0.6±0.1 embryo lengths; Figure 1I). This difference is most likely due to long-ranged flow requiring a certain degree of stable, filamentous network components. These kinetic differences also seem to contribute to NMY-2 accumulating at the furrow while F-actin does not accumulate at that site (Figure 1I). This is most striking during late cytokinesis where substantial amounts of linearly organized NMY-2 still flow towards the future midbody while F-actin does not show any recognizable flow at that stage (Figure 1F). These observations suggest – similar to what has been recently found in mammalian tissue culture [48] – that non-muscle myosin II might also be organized in aligned stacks in the *C. elegans* cortex that can span several micrometers and whose turnover is independent of the turnover of actin filaments.

### 3.2 Uniaxial loading counteracts longitudinal flows

In order to probe cytokinesis mechanics, we used the well-established parallel plate assay [35–37]. To achieve highly consistent uniaxial loading in the parallel plate assay, we employed monodisperse, inert beads with diameters of 25, 20, 15, and 13.5 μm, (representing 0, 20, 40, and 46% uniaxial compression, respectively; Video 2). Uniaxial loading induces a shape anisotropy where the surfaces contacting the plates become flat and the remaining surfaces start to bulge. Importantly, it has been shown that uniaxial loading directly impinges on cortex mechanics since (a) the cell boundary is governed by Laplace’s law (Figure 2A) [37]; (b) external friction (friction between the actomyosin cortex and the plasma membrane/vitelline membrane/egg shell) can be neglected [29, 49]; (c) the elastic cortical layer dominates cell mechanics in the system while the contribution of the plasma membrane can be largely ignored [38, 49, 50]. Analyzing longitudinal NMY-2 cortical flow prior to the onset of furrowing, we found that longitudinal flow velocities are highest in unloaded embryos and decrease with increased loading (Figure 2B, left). Flow velocities were down to 3.5 and 3.4 μm/min in anterior and posterior domains, respectively, in 20% compressed embryos and decrease further to 1.8 and 2.8 μm/min with 40% compression (Figure 2B; Figure S1A). Wt embryos compressed by 46% reach only −0.7 and 1.8 μm/min and fail to cleave (Video 2). The strong reduction of longitudinal flow (flow along the a-p axis) is best apparent in superimpositions of consecutive frames from time lapse recordings (Figure 2B, right). Interestingly, the reduction of longitudinal flow scales to the amount of loading, suggesting that the cortex behaves like an elastic material (Figure S1B).

**Figure 2.**
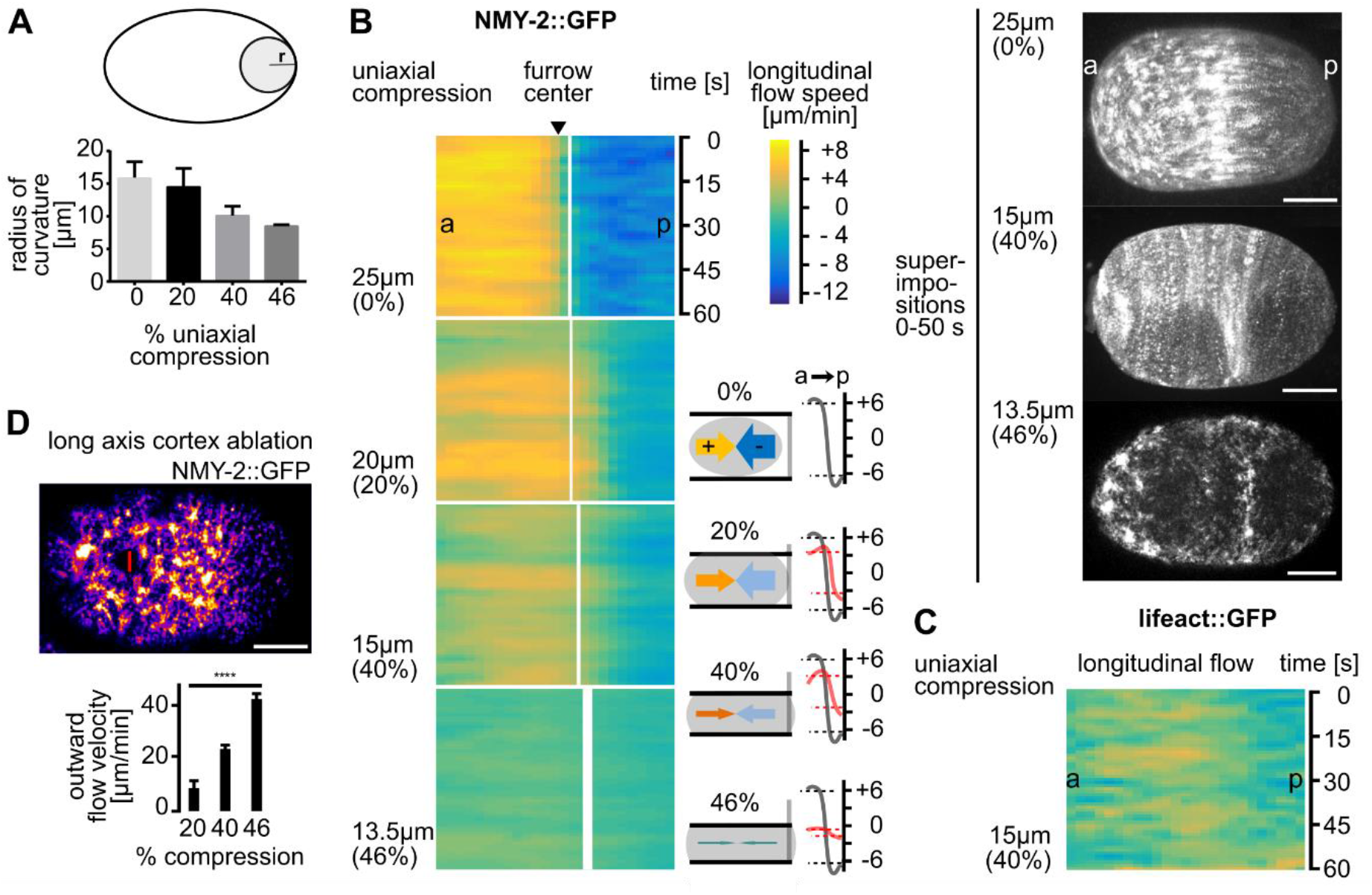
Longitudinal NMY-2 flow is mechanosensitive. **(A)** Quantification of curvature increase due to compression. Smaller radii represent higher curvature (see cartoon and Methods). **(B)** Left: Heat map kymographs of cortical flow velocities obtained from PIV of NMY-2::GFP foci moving along the long axis of differently mounted one-cell *C. elegans* embryos. For statistical parameters of heat maps see Fig. S1A. Black arrow head points to the white line demarcating the future furrow. Thickness of the line represents standard deviation. Bottom middle: Paradigm of uniaxial compression and corresponding flow velocities. Bottom right: Averaged velocities (over 60 s) along the anterior-posterior (a-p) axis from the PIV analysis (right panels). Grey and red lines represent averaged velocities in uncompressed and compressed embryos, respectively (n = 5 each). Right: Superimpositions generated by overlaying stills from projected time lapse images. Scale bars = 10 μm. **(C)** Heat map kymographs generated by PIV of lifeact::mCherry for longitudinal flow. Embryos were imaged under 40% compression (n = 5). **(D)** Top: Representative still from NMY-2::GFP expressing embryo exhibiting a cortical wound inflicted by UV laser cutting along the short axis of the embryo. Left: Quantification of outward flow velocities following cortical wounding under increasing compression (n = 5 each). See also Video 3.

Consistent with F-actin having faster cortical turnover, we find slightly weaker and less uniform longitudinal flow for F-actin (lifeact) compared to NMY-2 (Figure 2C, Figure S1A). Since uniaxial compression induces a shape anisotropy that leads to anisotropic stress in the cortex [38], this might alter cortical tension and impinge on longitudinal cortical flows. To test this, we performed cortical laser ablations [29] just prior to the onset of polarizing flow after fertilization parallel to the short axis of the embryo (cuts of 23% embryo width; Figure 2D, Video 3). We chose this time point for ablations since the cortex shows a highly similar architecture to the cortex just prior to cytokinesis [9] and the measurements are not confounded by fast changing patterns of flows. We made sure that the cortical wound induced by laser ablation did not vary in size under different degrees of compression (Figure S1C). Measuring outward velocities of NMY-2 foci post ablation, we found that increased loading generates increased outward flow velocities (11±0.6 μm/min at 20% compression, 23±1 μm/min at 40% compression, and 43±2 μm/min at 46% compression; Figure 2D). Notably, our ablation experiments only allow measurements of changes in total mechanical stress but not the relative contribution of passive and active stresses. Although our ablation experiments were performed before onset of cytokinetic flows, they clearly demonstrate a response of the cortex that scales to loading nevertheless. Thus, our observations are consistent with the interpretation that uniaxial compression induces cortical stress which seems to counteract longitudinal flows (Figure 2B) and eventually prevents successful furrowing.

### 3.3 Rotational flow is induced upon uniaxial loading

Work from our lab and others has uncovered rotational flow of the cortex – which is orthogonal to longitudinal flow – in the one-cell *C. elegans* embryo directly before contractile ring formation (Figure 3A, top left) [19, 20]. This rotational flow also occurs *in utero* (Figure 3A, left) and is most likely due to deformations of embryos similar to 20-40% uniaxial loading when measuring circularity of embryos in utero and comparing this to contact angles measured ex utero for uncompressed embryos (Figure 3A, right). However, the questions whether this flow is an intrinsic property or whether it needs a trigger and how it contributes to cytokinesis itself have not been addressed so far. Utilizing the paradigm of the uniaxial loading by the parallel plate assay, we observed that while longitudinal NMY-2 flow velocities decrease, rotational cortical flow velocities increase concomitantly (Figure 3B, Figure S1D, Videos 2,4), from 0.8±0.02 μm/min in uncompressed to a maximum of 23±0.1 μm/min in 40% compressed embryos. Under very high loading, rotational flow is virtually absent due to accumulation of NMY-2, F-actin and activated RhoA on bulging surfaces (see below). Remarkably, this shows that rotational flow is strongly enhanced by mechanical stress. Again, consistent with F-actin showing faster cortical turnover, we also find that F-actin shows a shorter range of rotational flow (Figure 3C, Figure S1D). More importantly, the magnitude of rotational cortical flow scales to the amount of loading (Figure S1E). Together with the scaling of longitudinal flows (Figure S1B), this strongly suggests that the two phenomena are not simply occurring coincidentally but that they are most likely interdependent.

**Figure 3.**
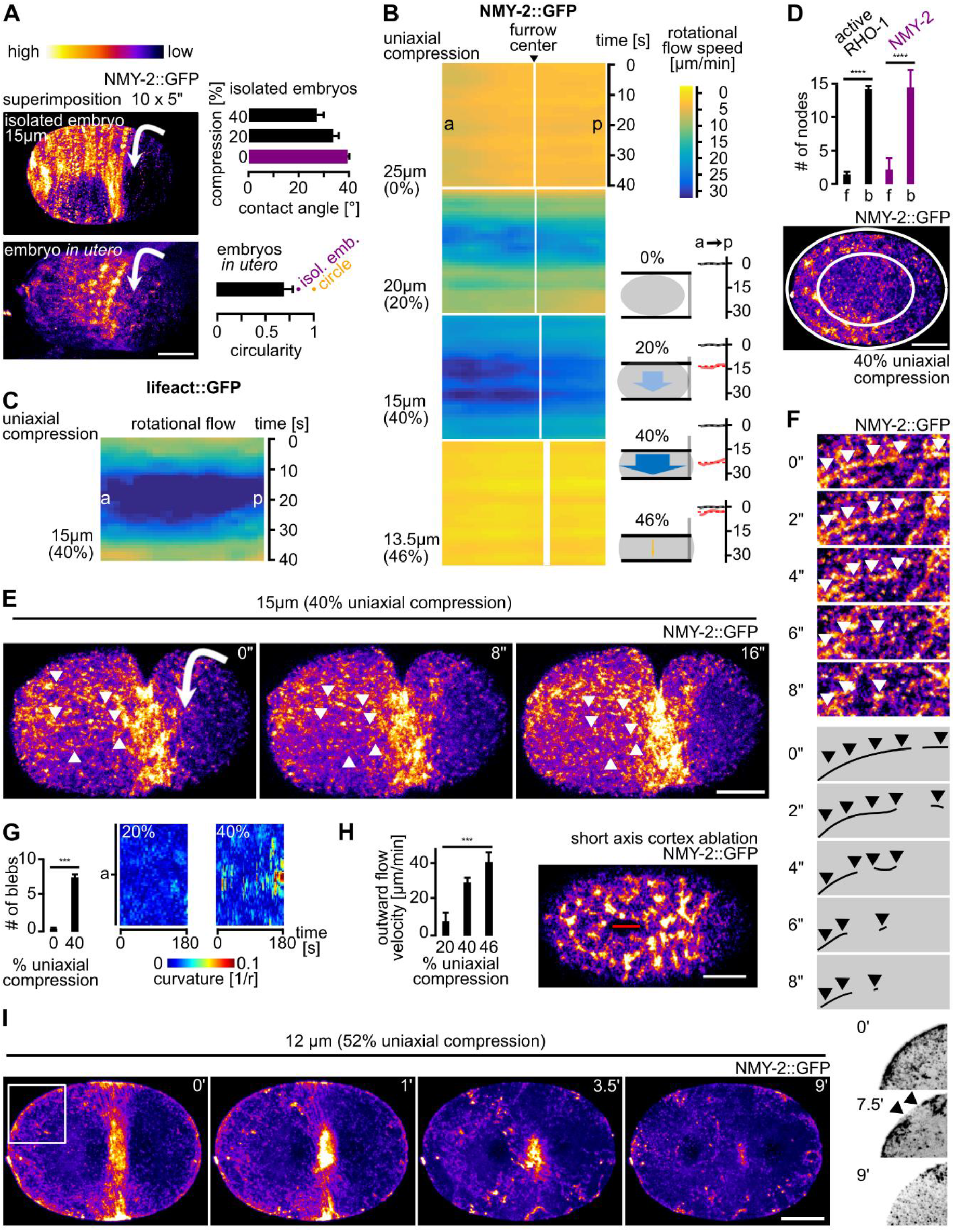
Rotational cortical flow is required for furrowing under uniaxial compression. **(A)** Left: Maximum projected stills from time lapse microscopy of a representative, isolated wt embryo (top) and an embryo inside the uterus (bottom); scale bar = 10 μm. Direction of cortical rotation is indicated by an arrow. Top right: Contact angles between coverslip and embryo. Bottom right: Circularity of embryos *in utero* (n = 6), circularity for ellipsoidal, isolated embryos and a circle are also included. **(B)** Heat map kymographs of cortical flow velocity values from NMY-2::GFP particle tracking along the short axis of differently mounted embryos. For statistical parameters of heat maps see Fig. S1D. Black arrow head points to the future furrow center. Bottom middle: Cartoon depictions of corresponding rotational cortical flow velocities. Bottom right: Averaged velocities (over 60 s) along the a-p axis from the PIV analysis (left panels). Grey and red lines represent averaged velocities in uncompressed and compressed embryos, respectively (n = 5 each). **(C)** Heat map kymographs generated by PIV of lifeact::mCherry for rotational flow. Embryos were imaged under 40% compression (n = 5). **(D)** Top: Quantification of active RHO-1 (black) and NMY-2 (purple) nodes on flat (f) versus bulging (b) surfaces in embryos under 40% compression (n = 5 each). Right: Representative projection of an embryo illustrating the quantification for NMY-2::GFP (inner ellipse = flattened surface; see panel E for fluorescence intensity color code). Scale bar = 10 μm. **(E)** Projections from time-lapse data (see Video 5). Arrowheads point to linear cortical NMY-2 that undergoes rupture. Scale bar = 10 μm. **(F)** Magnified projection of the cortex showing rupture of linearly organized NMY-2. Scale bar = 2.5 μm. **(G)** Left: Quantification of the number of blebs in uncompressed and 40% compressed WT embryos over 60 s. Right: Quantification of curvature changes. Two representative curvature kymographs for a 20% and a 40% compressed embryo are shown. See experimental procedures for details. **(H)** Left: Quantification of outward flow velocities following cortical wounding under increasing loads (n = 5 each). Right: Representative still from a NMY-2::GFP expressing embryo exhibiting a cortical wound inflicted along the long axis of the embryo by UV laser cutting. Scale bar = 10 μm. See also Video 6. **(I)** Cortex rupture for 52% compression. Representative projections from time-lapse microscopy are shown; scale bar = 10 μm. The right pictures show the boxed area of the leftmost still annotated with arrowheads and inverted to illustrate cortex rupture. See also Video 8.

Based on these findings we asked how stress created by uniaxial compression [38] contributes to rotational cortical flow. Analyzing the distribution of NMY-2, F-actin and active RHO-1 (using a RhoA sensor consisting of GFP fused to the AH-and PH-domains of ANI-1; [31]) we found cytokinetic nodes assembling uniformly in uncompressed embryos. In contrast, in compressed embryos, NMY-2, F-actin, and active RhoA are only found at the equator and on bulging surfaces (Figure 3D), for which it has been shown that their cortex is more stressed [4, 7, 38]. This suggests that cell cycle-dependent RhoA activation can be local and most likely in response to cortical deformation. Shortly after their assembly, focally and linearly organized NMY-2 moves onto flattened surfaces through rotational flow (Figure 3E; Video 5). Due to actomyosin being concentrated on bulging surfaces in loaded embryos, its mobilization by rotational flow generates a flow front – the former boundary between the bulged and flat cortex – that moves over the flattened surface until the front reaches the bulged surface on the other side (Video 5).

Moreover, while linearly organized NMY-2 connecting cytokinetic nodes in uncompressed embryos constricts, it ruptures in compressed embryos (Figure 3E, 3F; Video 5). Rupture occurs anisotropically in the direction of rotation, starting at the front of rotational flow (Video 5). Notably, this anisotropy correlates with the asymmetric position of the midbody, midbodies always forming where rotational flow emerged, opposite to the side where cortex filament rupture occurs at the rotational flow front (Video 5). This always leads to asymmetric positioning of the midbody (n>20; data not shown). Additionally, rupture leads to both flow towards the furrow (from the furrow-facing side of the rupture) and flow towards the poles (from the pole-facing side of the rupture) (Video 5). Flows towards the furrow have similar velocities as longitudinal flows in uncompressed embryos and can lead to similar parallel alignment of cortical material in the equatorial zone (Figure 1C, 1E). Flows towards the poles dissipate due to dissolution of nodes and lack of a barrier similar to the equatorial band of focal and linear NMY-2 (Video 5). Furthermore, these flows occur at the same time as polar blebbing is observed, which might additionally contribute to cortical relaxation of cortical tension caused by pole-directed cortical flow (Figure 3G).

Since uniaxial compression leads to anisotropic cortex assembly at the onset of cytokinesis and anisotropic disassembly during furrowing, we asked whether loading induces anisotropies in cortical tension that could also contribute to rotational flow. To test this, we performed laser cutting of the cortex (cuts of 16% embryo length; Figure 3H, Video 6) parallel to the long axis of the embryo just prior to the onset of polarizing flow after fertilization and observed a loading-dependent increase in initial outward flow velocities of NMY-2 particles at the site of the cortical wound (15±0.5 μm/min at 20% compression, 29±2 μm/min at 40% compression, and 32±4 μm/min at 46% compression; Figure 3H).

When measuring outward velocities 5 s after cortex ablation (as established previously; [29]), it seems that tension increases along the short axis scales more linearly with loading (Figure 3H, R^2^ = 0.94, Figure S1G) than along the long axis (Figure 2D; R^2^ = 0.83, Figure S1G). Also consistent with previous work [29], tension seems to be higher along the short axis under low loading. Given the elegant theoretical framework of cortical mechanics that highlights the roles of effective viscosity and local compression rate for the generation of polarizing cortical flow [29], the above measurements suggest that besides cortical stress, viscosity and/or cortex compressibility might additionally contribute to rotational flow.

### 3.4 Uniaxial loading and the limit of cytokinetic mechanostability

Next, we asked how rotational flow changes when we subject embryos to 46% compression, a load where embryos do not divide (Video 7). Here, we found the same anisotropic distribution of nodes as for 20% and 40% compression, however, nodes on bulged surfaces do not translocate by rotational flow. Instead, streaming of linearly organized NMY-2 in the equatorial area is observed (Video 7). Streaming does not lead to the bundling of linear NMY-2 at the equator and a contractile ring is not formed (0% of embryos; n>15). Moreover, under 46% compression, actomyosin recruitment to the equatorial zone by the central spindle pathway can still be observed, however, equatorial actomyosin recruitment is insufficient for furrowing. Similar to human cells [37], we found that the limit of cortex loading is reached at 52% (12 μm beads; 50% for human cells). Due to increased bulging, the cortex ruptures at these bulged sites and the equatorial NMY-2 band disintegrates (Figure 3I; Video 8). This confirms that the cortex is bearing the load of compression since we neither observed rupture of the plasma membrane nor of the eggshell. Moreover, it also supports the idea that the cortex behaves like an elastic material that has a yield point at 52% compression.

### 3.5 Actin-myosin regulators required for mechanostable cytokinesis

Since cortical tension along the short axis is under the control of the Rho GTPase cycle (Meyer et al., 2010) and since NMY-2 cortical polarization is observed independently of the level of loading, we performed a targeted screen to identify factors involved in cortical rotation and linear organization. For the screen, we used 20% and 40% compression of embryos, where strong rotational flow is observed in wild-type embryos (Figure 4A). This screen identified several factors including (1) MEL-11, a myosin-associated phosphatase [51], required for both focal and linearly organized NMY-2 (Figure 4B; Video 9); (2) LIN-5, a factor known to regulate spindle positioning [52], which also promotes the transition from focal to linear organization and seems to stabilize the latter (Figure 4C; Video 10); (3) ECT-2, a cytokinesis regulatory RhoGEF [53], which is required for proper size and density of focal and linear NMY-2 (Figure 4D; Video 11); (4) RGA-3, a cytokinesis regulatory RhoGAP [54, 55], for which it has been previously shown that depletion leads to exaggerated rotational flow [44], and which we find is also required for node formation and to suppress excess linear organization (Figure 4E; Video 12); (5) NOP-1, a factor required in parallel with the RhoGAP CYK-4 to promote RHO-1 activation and NMY-2 node formation during cytokinesis [31], which is also required for the transition to linearly organized NMY-2 (Figure 4F; Video 13); and (6) POD-1, a type III Coronin implicated in actin dynamics and crosslinking [56], which is as well required for this transition (Figure 4G, top panels; Video 14). Moreover, *pod-1* RNAi leads to the formation of short-lived circular contractile NMY-2 structures, which suggests that Coronin-mediated actin crosslinking is required to coordinate formation of long-range NMY-2 linear organization to achieve pole-to-equator flow (Figure 4G, bottom panels; Video 14).

**Figure 4.**
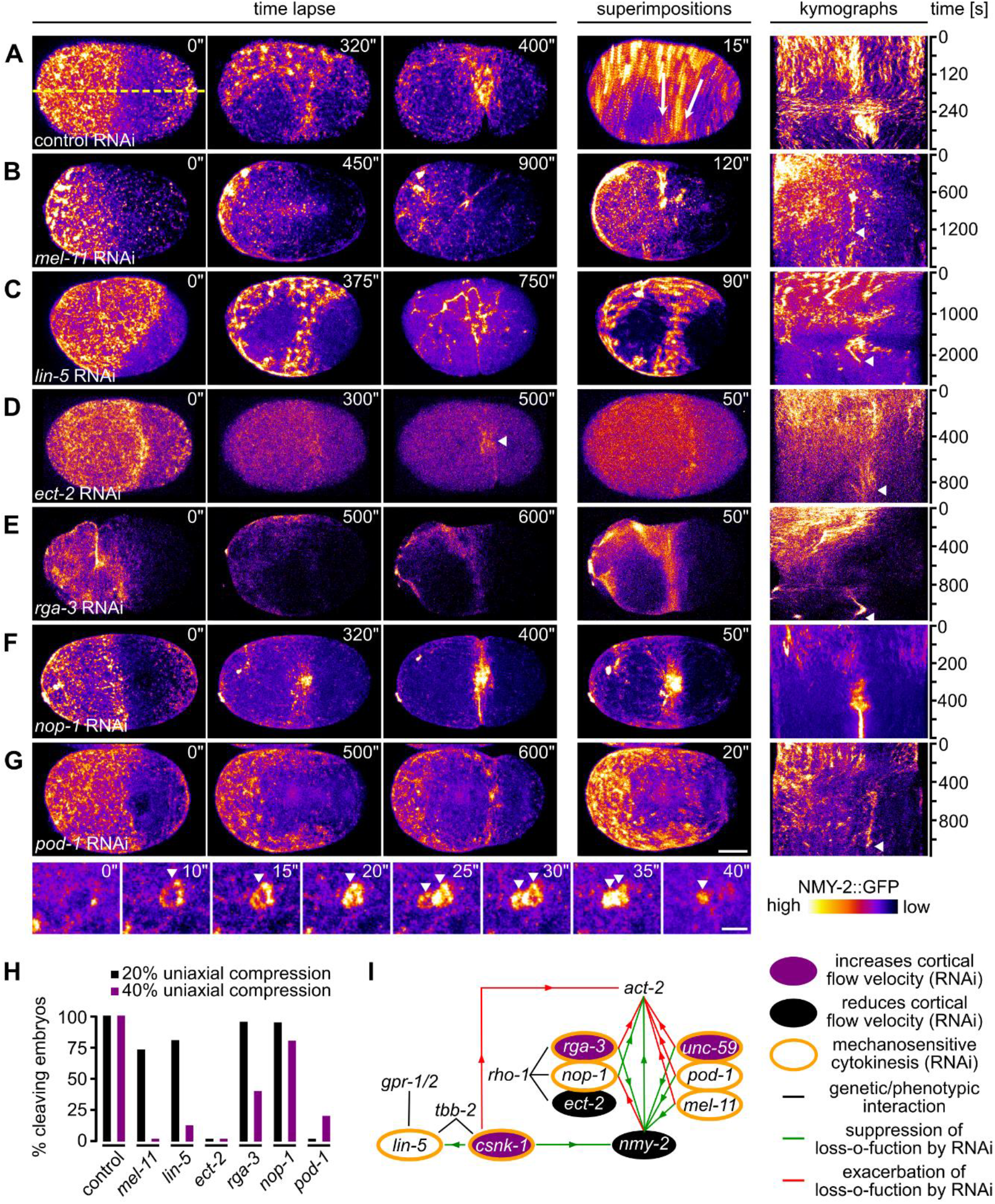
Antagonistic actin-myosin regulators are required for rotational flow and cytokinesis mechanostability. **(A)** Left: Maximum projected stills from time lapse microscopy of a representative wt embryo expressing NMY-2::GFP. Middle: Superimposition of frames from a 15 s time window. White arrows indicate direction of rotational flow. Right: Kymograph generated along the dashed yellow line in the leftmost panel. **(B-G)** Representations as in panel (A) but for embryos treated with the indicated RNAi. Scale bar = 10 μm. See also Videos 9-14. Bottom of panel (G): Magnification of projected stills showing formation of cortical circular structures (arrowheads) in *pod-1* RNAi embryos. Scale bar = 2.5 μm. See also Video 14. **(H)** Quantification of successful first cell division for the indicated RNAi treatments under 20% (black) and 40% (purple) compression (n≥5 each). **(I)** Genetic network of factors controlling cytokinesis. Interactions are based on [21, 31] and data from panel H.

Although RNAi of these regulators gives rise to very distinct phenotypes, for all factors where furrowing phenotypes were not known (MEL-11, LIN-5, RGA-3, NOP-1, and POD-1), we observed a loading-dependent failure of cytokinesis completion (Figure 4H). With the exception of *pod-1* RNAi, increased loading leads to an exacerbation of the phenotype. Remarkably, all regulators are known to have opposing phenotypes in actin (*act-2*) and myosin (*nmy-2*) mutants [32] and are directly or indirectly linked to the Rho GTPase cycle (Figure 4I). This network of factors is essential for cytokinesis’ mechanical robustness and by differentially regulating NMY-2 organization seems to indirectly also affect cortical viscosity and compressibility.)

### 3.6 Persistent linearly polarized NMY-2 prevents cortical rotation

Previously it was shown that *rga-3* RNAi leads to exaggerated chiral flows during a-p polarization of the one cell *C. elegans* embryo [21, 44]. However, the data above shows that *rga-3* RNAi embryos do not divide under uniaxial compression. We therefore more closely investigated the origin of exaggerated chiral flows in rga-3 RNAi embryos and why this prevents cytokinesis under mechanical stress. Although we observe the reported exaggerated chiral flow during a-p polarization under uniaxial loading (Figure 5A), an important additional phenotype of *rga-3* RNAi embryos is persistent and long range linearly organized cortical NMY-2, which can be observed both during a-p polarization (Fig. 5A) and right after the onset of cytokinesis (Figure 5B). This organization is maintained during cytokinesis and leads to peeling of the filaments towards the nascent midbody, lack of a proper contractile ring (Figure 5B), and strongly reduced rotational flow under load (Figure 5C). Thus, unlike in wt embryos, linearly organized cortical NMY-2 does not undergo remodelling or ruptures in *rga-3* RNAi embryos. Considering the theory of cortical torques [21] it seems likely that a persistent linear organization of NMY-2 can induce stronger and more long ranged torques than wt. We propose that this leads to excessive chiral flow during polarization but later results in lack of cortical rotation and failed cytokinesis long-range linear connections are not remodelled (Figure 5B).

**Figure 5.**
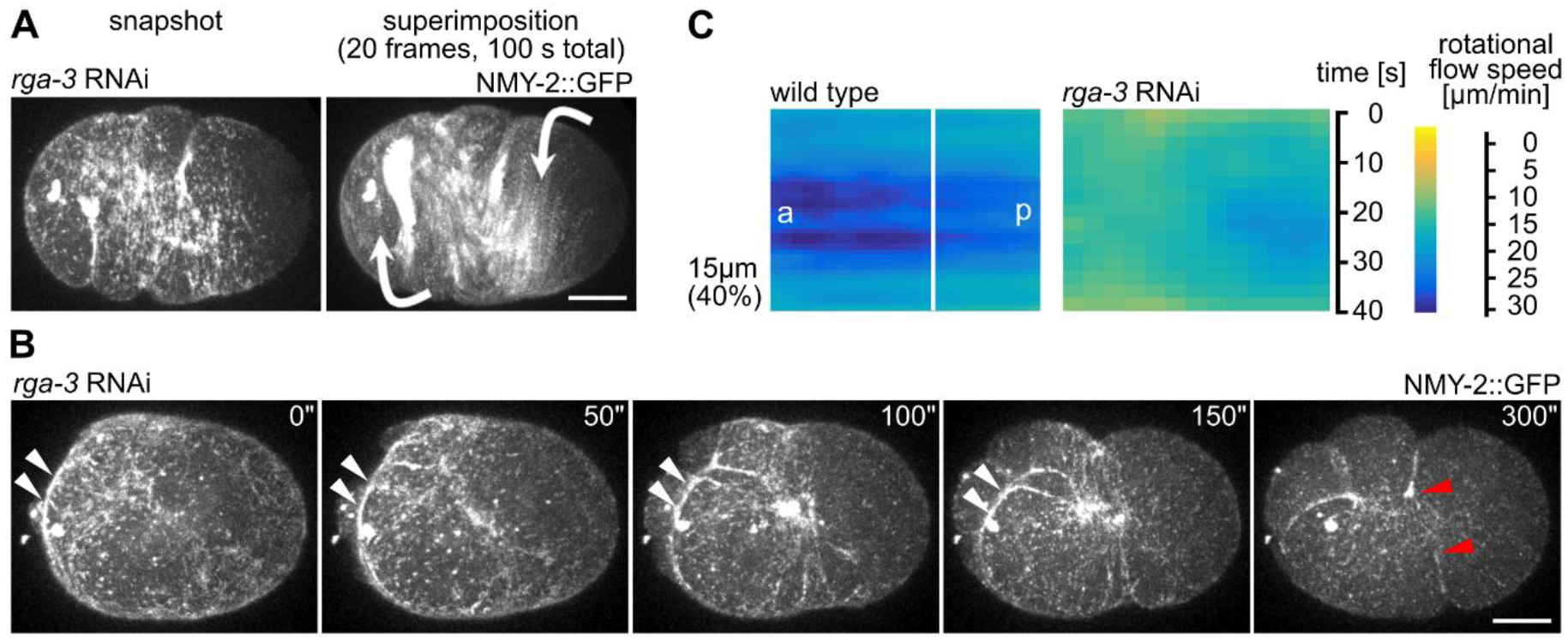
*rga-3* RNAi leads to increased linear organization of cortical NMY-2 and lack of rotational flow under load. **(A)** Still and superimposed stills from time-lapse microscopy of a representative *rga-3* RNAi embryo. Note the linear organization of NMY-2 and the almost exclusive rotational trajectories of cortical NMY-2 in the superimposition. Direction of rotational trajectories (arrows) have opposite polarity (anterior domain counterclockwise and posterior domain clockwise. Scale bar = 10 μm. **(B)** Stills from a time-lapse series of a representative *rga-3* RNAi embryo during cytokinesis. White arrowheads mark long linear cortical NMY-2 that peels from the sides towards the nascent midbody. Red arrowheads mark the dissolving furrow. Scale bar = 10 μm. **(C)** Heat map kymographs of rotational cortical flow velocity values from NMY-2::GFP particle tracking along the short axis of wt and *rga-3* RNAi embryos mounted under 40% uniaxial compression (n =5 each).

### 3.7 Cortical chirality and polarity are required for rotational flow polarization

Based on previous findings demonstrating that the actomyosin cortex generates active chiral torques with invariant handedness important for axial patterning [19–21], we reasoned that regulators of cortical chirality will contribute to rotational cortical flow polarization. To do so, we used RNAi inhibiting the expression of the casein kinase 1γ, CSNK-1. In line with earlier observations [20], we found that in *csnk-1* RNAi embryos, rotational cortical flows can switch their handedness across the equator and, concomitantly, a strong reduction of compressivelongitudinal flow occurs (Figure 6A, left; Figure S2A). Importantly, the switch of rotational flow handedness generates shear flow in the equatorial region, which leads to dissolution of the furrow under mechanical load (Figure 6A, right, 40% compression; Video 15). This phenotype is not restricted to *csnk-1* RNAi embryos, but also occurs when components of the Wnt pathway that have been shown to be required for cortical torque generation and chiral symmetry breaking are targeted by RNAi [21, 39], for instance *mom-2* (Figure 6B).

**Figure 6.**
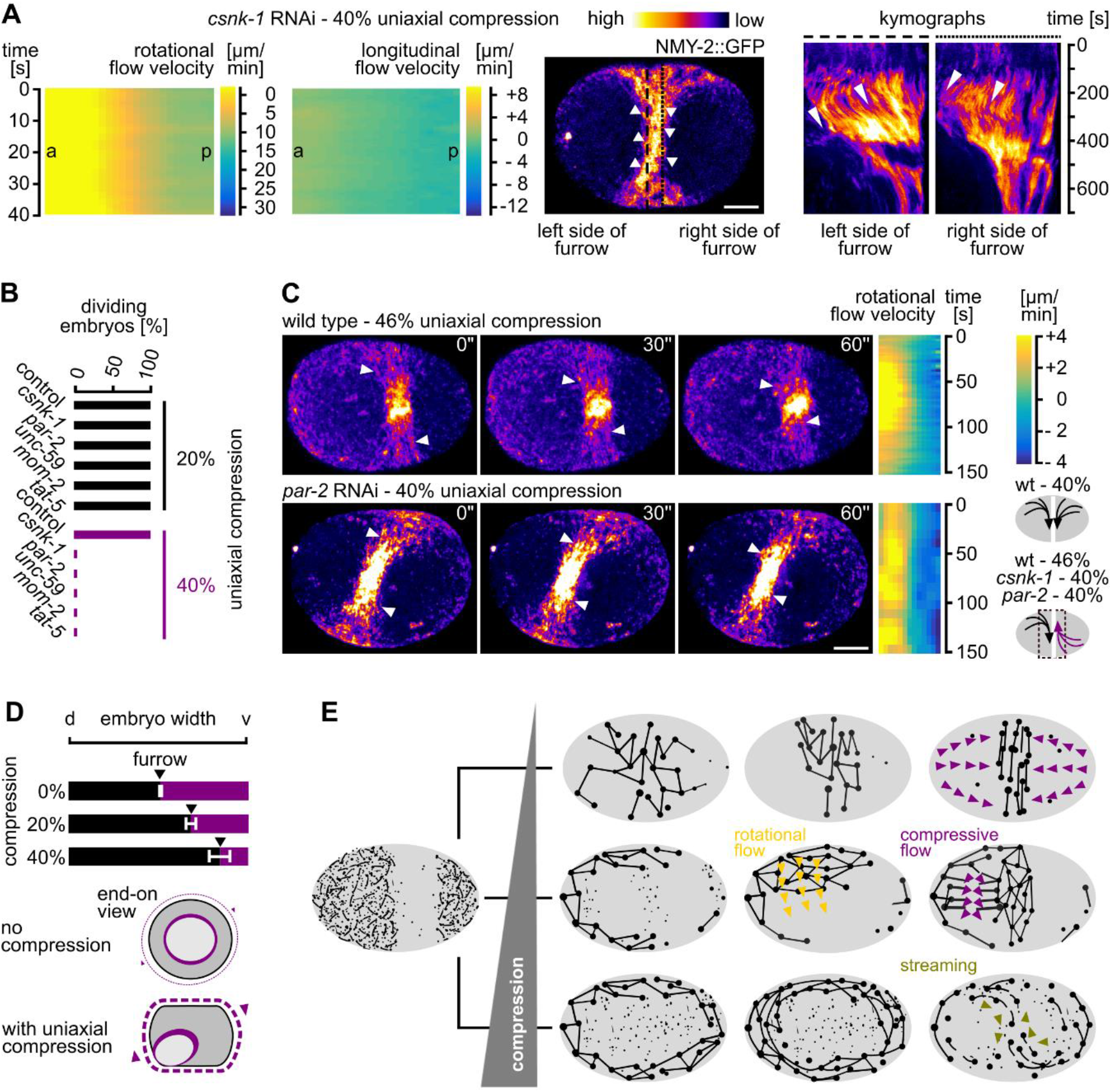
Cortical polarity and chirality are required for mechanostable cytokinesis. **(A)** Left: Heat map kymographs generated by PIV of NMY-2 particles along the short and the long axis of one-cell *C. elegans csnk-1* RNAi embryos. All embryos were imaged under 40% compression (n = 5 each). Middle: Maximum projected still from time lapse microscopy of a representative *csnk-1* RNAi embryo expressing NMY-2::GFP; white arrowheads indicate flow direction in the furrow region; scale bar = 10 μm. See also Video 15. Right: Kymographs generated along the dashed lines at the left and right boundary of the furrow. Opposite polarity of flow is indicated by arrowheads. Note the dissolution of the furrow after 400 s. **(B)** Quantification of successful first cell division for the indicated RNAi treatments under 20% (black bars) and 40% (purple bars) compression (n = 10 each). **(C)** Left: Maximum projected stills from time lapse microscopy of a representative wt (46% compressed) and *par-2* RNAi embryo (40% compressed) expressing NMY-2::GFP; scale bar = 10 μm. See also Video 16. Middle: Heat maps generated via PIV of NMY-2 particle flow in the furrow region along the short axis. Bottom right: Cartoons depicting rotational flow polarization in wt (top) and RNAi embryos (bottom). **(D)** Furrowing asymmetry quantified in wt embryos. Top: Average furrow position along the short axis is indicated by black arrowheads (n = 5 each). Bottom: Model how lack of rotational cortical flow influences furrow asymmetry. See text for details. **(E)** Model for linearly organized cortical myosin dynamics under different conditions. Left: Cortical NMY-2 distribution before the onset of cytokinesis. Top right: Linear and focal NMY-2 coalesce into an equatorial band in unstressed embryos through longitudinal pole-to-equator flow (purple arrowheads). Middle right: With increased loading, NMY-2 foci show an anisotropic distribution at the onset of cytokinesis, subsequently, focal and linear NMY-2 show rotational flow (orange arrowheads) and linearly organized NMY-2 ruptures, thereby generating longitudinal flow (purple arrowheads). Bottom right: With high load, anisotropically distributed nodes transform into a linearly organized network that shows streaming, preferentially in the equatorial region.

Since the contractile ring forms by alignment of linearly organized cortical materialfilaments through RhoA-dependent flow, the whole system also needs to be polarized along the direction of longitudinal compressive flow. Accordingly, we find that disruption of anterior-posterior polarity in *par-2* or *tat-5* RNAi embryos phenocopies the *csnk-1* and *mom-2* RNAi, a lack of longitudinally polarized compressive flow and shear flow in the equatorial region (Figure 5C, Figure S2B, S2C; Video 16). Importantly, we also observed shear flow in wt embryos when they are compressed by 46% and do not divide (Figure 6C; Video 7). This suggests that under these conditions uniform rotational cortical polarization that is observed in wt embryos up to 40% compression fails after removing factors responsible for cortical polarity and chirality or by excessive loading (Figure 6C, bottom right). Next, we asked how furrowing itself is affected by uniaxial loading and we tested whether factors known to be required for the intrinsic asymmetry of furrowing such as *unc-59* (encoding a septin; [27]) are also involved (Figure 6B). Similar to the requirement of genes involved in cortical polarity and chirality, we also found that *unc-59* RNAi embryos lack rotational cortical flow (data not shown) and fail to divide under 40% compression (Figure 6B).

Taken together, although factors involved in cortex polarity, chirality and asymmetry have not been found to be essential for cytokinesis in previous studies, they all become essential for cytokinesis under mechanical stress (Figure 6B, 6C). Furthermore, since compression induces rotational flow and all of the above RNAi embryos also show a loss of uniform polarized rotational flow (Videos 15, 16), we measured the degree of asymmetric furrowing under increasing mechanical load. In accordance with the above findings, we found that furrowing becomes increasingly asymmetric with increased loading (Figure 6D, top). These results, although correlative, strongly suggest that loading-induced rotational flows are involved in symmetry breaking during furrowing (Figure 6D, bottom).

## 4. Discussion

Our data outline a poorly uncharacterized feature of cortical flow, its mechanosensitivity and – up to a certain stress level – its mechanostability due to its ability to re-polarize from longitudinal to rotational (Figure 6D). Moreover, we demonstrate that uniaxial compression is a straightforward experimental paradigm to systematically investigate the mechanobiology of cortical flow during asymmetric cell division. Importantly, this paradigm shows that the induction of rotational flow depends on the magnitude of total mechanical stress. We also show that re-polarization of cortical flow is followed by anisotropic cortex rupture (Figure 6D). Rupture can lead to equator-directed cortical flows during cytokinesis which result in cortical compression around the cell equator and furrowing. This seems to be one mechanism that can balance extrinsic and intrinsic forces during cytokinesis (Figure 2B, Figure 3B). These results therefore extend previous work that identified longitudinal flows as non-essential contributors to contractile ring formation [9, 17]. In addition, our results reveal that besides polarization of actin filaments through flow-alignment coupling [9], cortical non-muscle myosin II also shows flow-alignment coupling, however, by having much longer lifetimes, cortical NMY-2 shows higher flow velocities than F-actin and accumulates at the equator and in the midbody – unlike F-actin (Figure 1). The recent thorough characterization of long, linearly organized non-muscle myosin II stacks whose lifetime is independent of the neighboring F-actin filaments [48] together with our observation of different cortical flow profiles for NMY-2 and F-actin (lifeact) strongly suggests that non-muscle myosin II has roles during cell division that are separable from those F-actin, in particular during final stages of contractile ring constriction and midbody formation (Figure 1G, 1I). Moreover, the proposed attractive interactions between linearly organized non-muscle myosin II [48] might also explain why we observe NMY-2 flows with longer duration and range than F-actin flows.

Previously, it has been demonstrated that the actomyosin cortex of embryos can be viewed as an excitable medium. In such a medium cortical flow in the form of waves is observed due to rapid local auto-activation of RhoA at wave fronts and delayed F-actin-mediated RhoA inhibition at the back of waves [57]. Treating the actomyosin cortex as an excitable material can therefore explain how the spindle determines the site of cleavage during cytokinesis, namely by generating signals that tune the auto-activation/inhibition cycle [57], it might, however, also explain the phenomena observed in this study, namely local activation of RhoA that leads to locally restricted non-muscle myosin II activation and stress-dependent flow polarization. In the framework of an excitable material, it seems most likely that local RhoA activation in uniaxially loaded *C. elegans* embryos is due to cell cycle state-dependent local auto-activation of RhoA on bulged areas of the cortex through a previously characterized mechanosensitive positive feedback [2]. The spatially restricted RhoA activation at these sites would then lead to the formation of a flow front during rotational flow (Video 5). Moreover, it also seems likely that anisotropies in spindle organization and spindle-cortex contacts pattern local auto-activation and thereby flow polarization. Thus, it is tempting to speculate that strong cortical flows are restricted to cytokinesis since the cortex only shows sufficient excitability during this specific cell cycle state and that only during cytokinesis the spindle or external mechanical forces can induce patterned activation/inactivation of RhoA that will generate polarized flows.

In addition, we demonstrate that several pathways which all have specific, non-redundant functions outside cytokinesis, fulfil essential roles for rotational cortical flow and furrow stability when cells are mechanically stressed (Figures 4, 5, 6). These pathways include the PAR and the Wnt pathway, which are known for their role in specifying the anteroposterior and the left/right body axes, respectively. Only for the PAR pathway a connection to cortical dynamics during cytokinesis is known (Jordan et al., 2016). Remarkably, interference with any of these pathways results in a similar mechanical stress-dependent failure of cytokinesis, a loss of uniform rotational cortical flow polarization, which leads to shear flow and dissolution of the contractile ring (Figure 6). This suggests that proper anteroposterior cortical polarization (*csnk-1, par-2, tat-5*) and yet to be identified aspects of cortical polarity that relate to left/right symmetry breaking or cortical torque generation (*rga-3, mom-2*; [20, 21, 39]) become essential for furrowing under mechanical stress. Additionally, we find that proper actomyosin regulation required for intrinsically asymmetric furrowing *(unc-59)* is also essential for cytokinesis mechanostability. This data supports earlier findings based on which it was argued that when the intrinsic asymmetry is disrupted, cytokinesis becomes sensitive to partial inhibition of contractility [27]. It should be noted that not only *csnk-1, rga-3*, and *unc-59* but also *par-2* and *tat-5* RNAi influence cortical cytoskeletal dynamics directly and further work involving super-resolution microscopy will be required to identify the origin of cortical cytoskeleton polarization during cytokinesis.

Although the data that we present here is correlative in many aspects, it nevertheless suggests that cortical rotation and cytokinesis mechanostability are intricately linked and rely on factors presumably required for symmetry breaking during cytokinesis, those that provide polarity information parallel (*csnk-1, par-2, tat-5*) and orthogonal (*rga-3, mom-2*) to the contractile ring and factors that potentially translate such polar bias into directional movement of actomyosin (*unc-59*). Moreover, our data also suggests that generation of cortical torque seems to depend on linear organization of cortical non-muscle myosin II (Figure 5). However, increased cortical torque alone is not sufficient for cytokinesis to proceed normally under load. Under these conditions, the remodeling of linear cortical structures seems crucial for the re-distribution of contractile cortical material towards the cleavage furrow by longitudinal flow and assembly of a contractile equatorial ring. Taken together, our findings show that Ray Rappaport’s notion that the cytokinesis machinery is ‘overbuilt, inefficient, never-failed, and repaired by simple measures’ [1] – in other words that cytokinesis is a robust process due to redundant regulators – might only be appropriate for unstressed cells, however, apparently redundant factors can become essential under mechanical stress.

## Author Contributions

Conceptualization, C.P.; methodology, D.S., O.D.; software, O.D.; validation, D.S., C.P.; formal analysis, D.S., O.D., and C.P.; investigation, D.S., C.P.; writing—original draft preparation, C.P.; writing— review and editing, C.P.; visualization, C.P.; supervision, C.P.; project administration, C.P.; funding acquisition, C.P.

